## Supplementary material for "Optimized tight binding between the S1 segment and KCNE3 is required for the constitutively open nature of the KCNQ1-KCNE3 channel complex": Figure supplements 1-17 & Table supplements 1-5

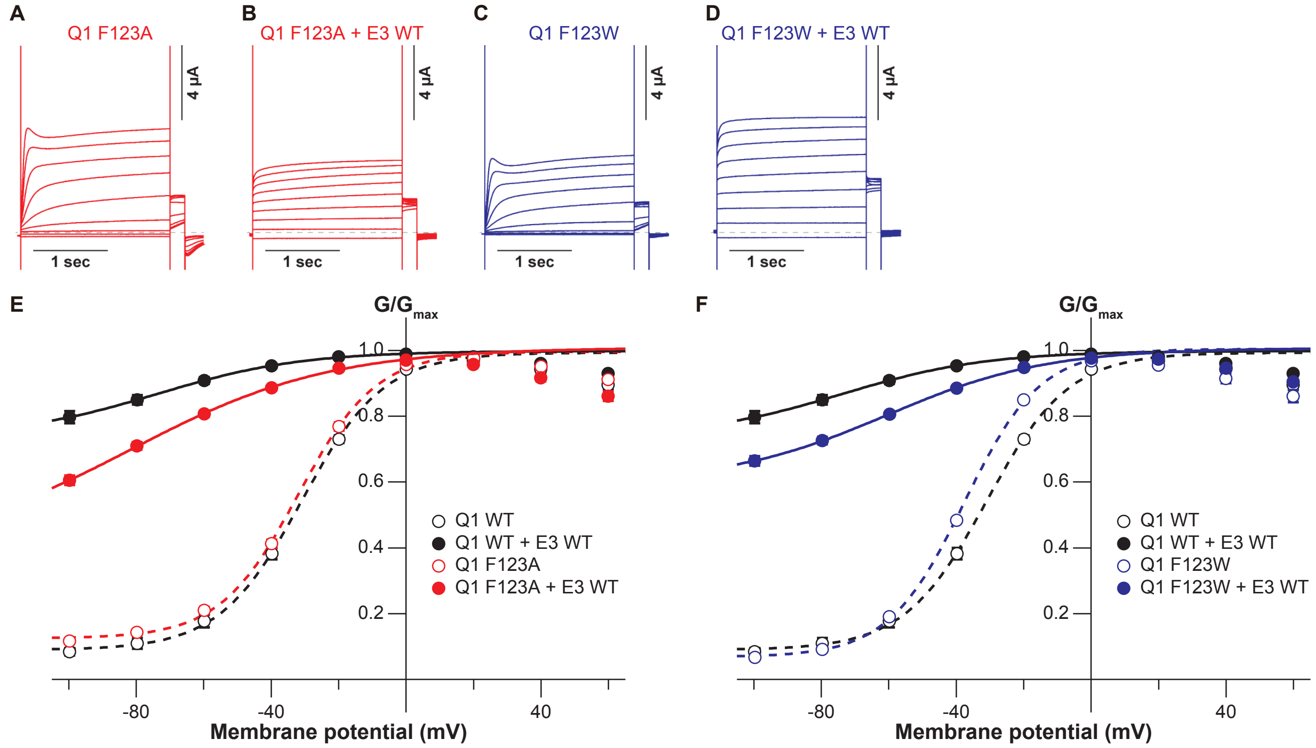

**Figure supplement 1, related to Figure 2. Current traces and G-V relationships of KCNQ1 F123 mutants.**

(**A-F**) Representative current traces (A-D) and G-V relationships (E,F) of KCNQ1 F123 mutants with or without KCNE3 WT. Error bars indicate ± s.e.m. for $n$ = 10 in (E,F).

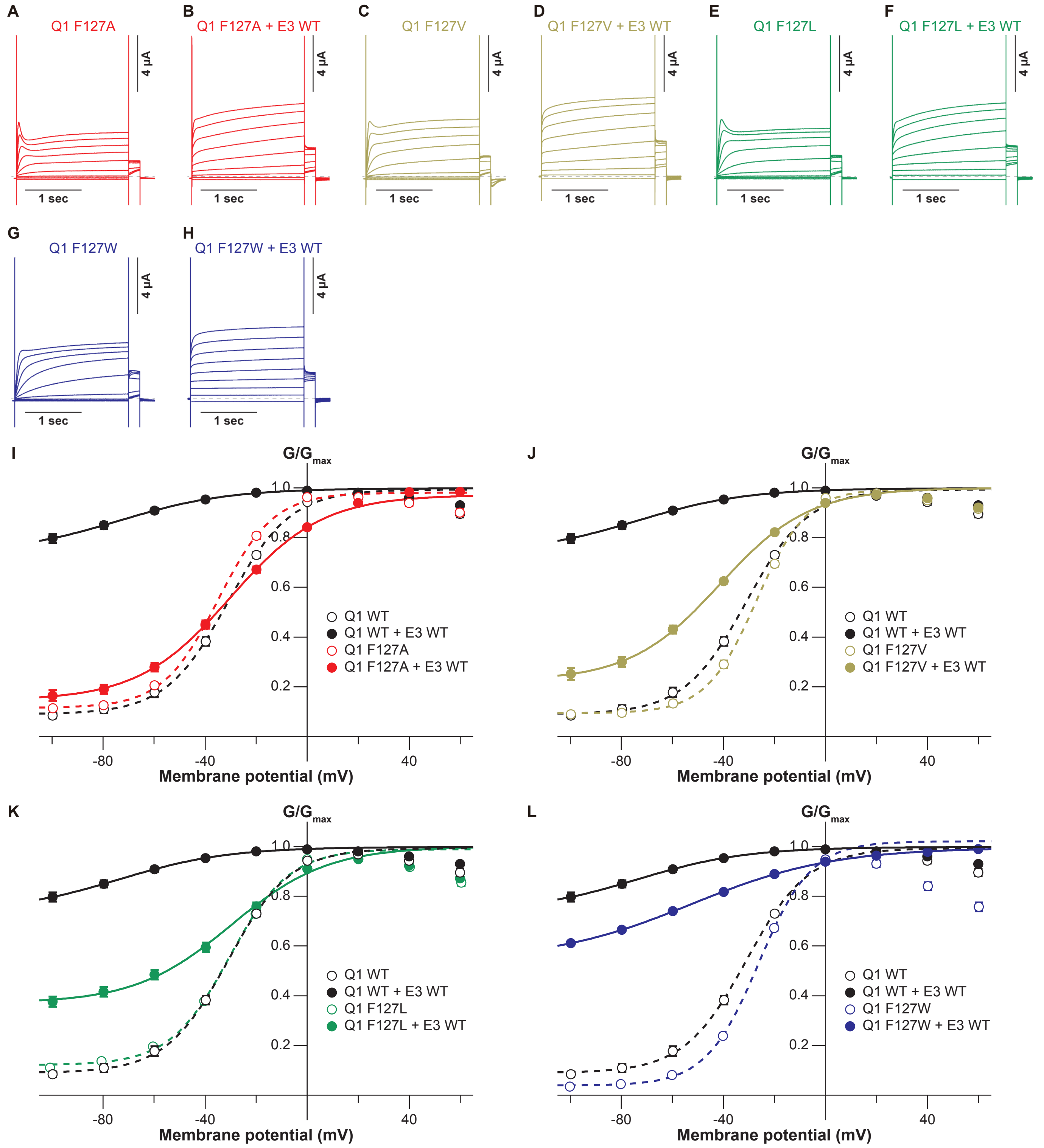

**Figure supplement 2, related to Figure 2. Current traces and G-V relationships of KCNQ1 F127 mutants.**

(**A-L**) Representative current traces (A-H) and G-V relationships (I-L) of KCNQ1 F127 mutants with or without KCNE3 WT. Error bars indicate ± s.e.m. for $n$ = 10 in (I-L). For clear comparisons, the current traces of KCNQ1 F127 mutants with KCNE3 WT shown in Figure 2C-F are redisplayed here.

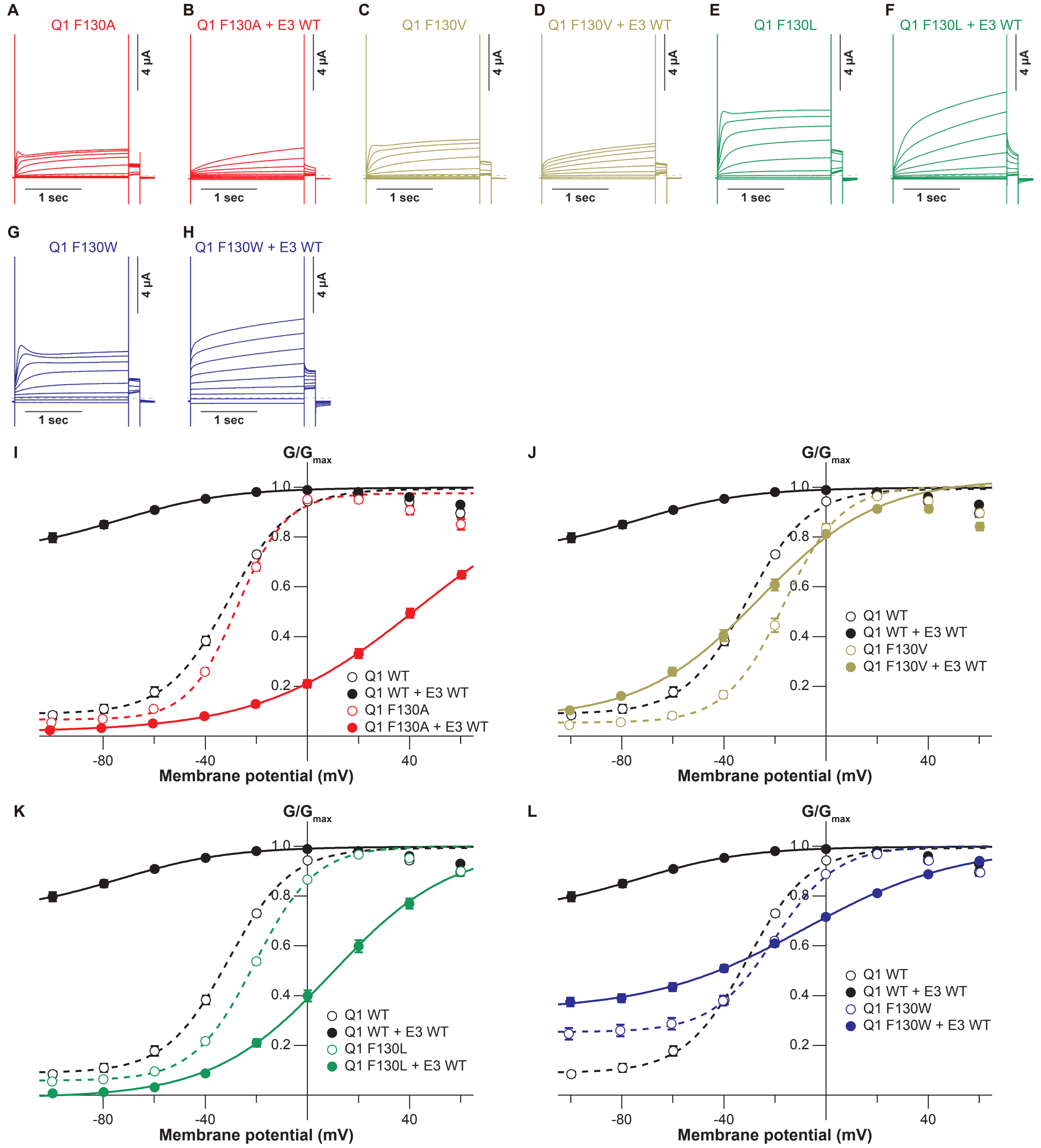

**Figure supplement 3, related to Figure 2. Current traces and G-V relationships of KCNQ1 F130 mutants.**

(**A-L)** Representative current traces (A-H) and G-V relationships (I-L) of KCNQ1 F130 mutants with or without KCNE3 WT. Error bars indicate ± s.e.m. for *n* = 10 in (I-L).

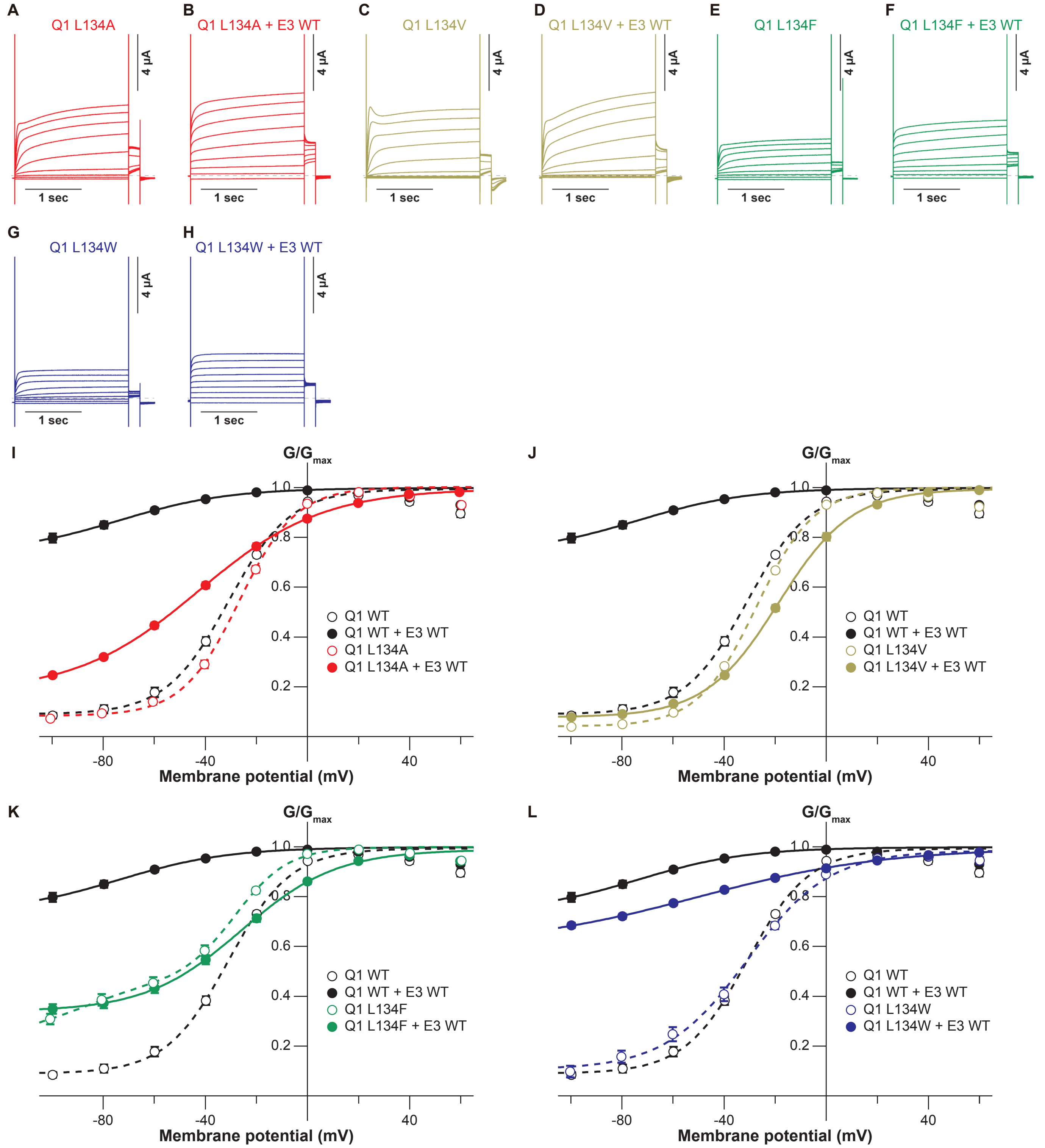

**Figure supplement 4, related to Figure 2. Current traces and G-V relationships of KCNQ1 L134 mutants.**

(**A-L**) Representative current traces (A-H) and G-V relationships (I-L) of KCNQ1 L134 mutants with or without KCNE3 WT. Error bars indicate ± s.e.m. for $n$ = 10 in (I-L).

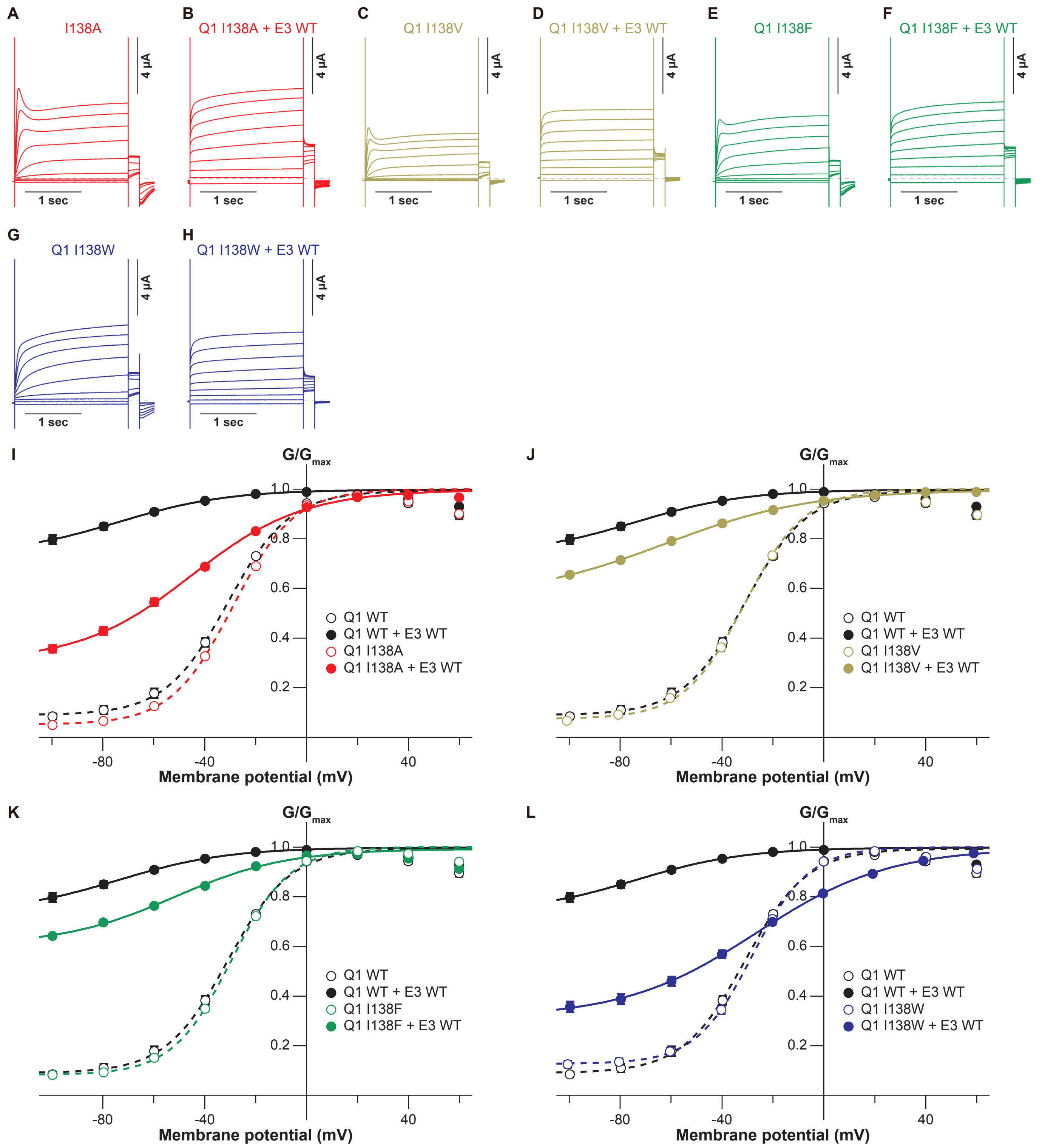

**Figure supplement 5, related to Figure 2. Current traces and G-V relationships of KCNQ1 I138 mutants.**

**(A-L)** Representative current traces (A-H) and G-V relationships (I-L) of KCNQ1 I138 mutants with or without KCNE3 WT. Error bars indicate ± s.e.m. for *n* = 10 in (I-L).

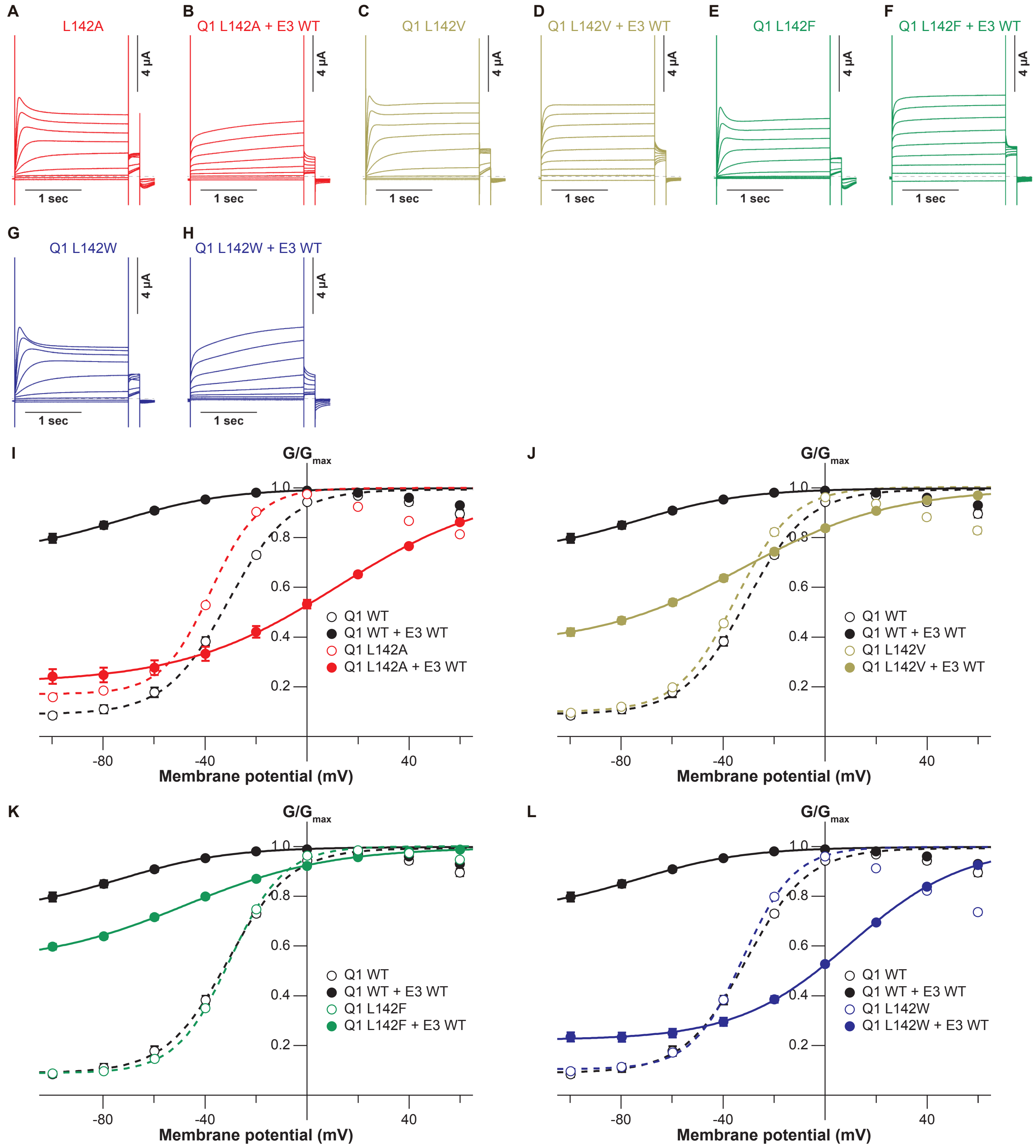

**Figure supplement 6, related to Figure 2. Current traces and G-V relationships of KCNQ1 L142 mutants.**

**(A-L)** Representative current traces (A-H) and G-V relationships (I-L) of KCNQ1 L142 mutants with or without KCNE3 WT. Error bars indicate ± s.e.m. for *n* = 10 in (I-L).

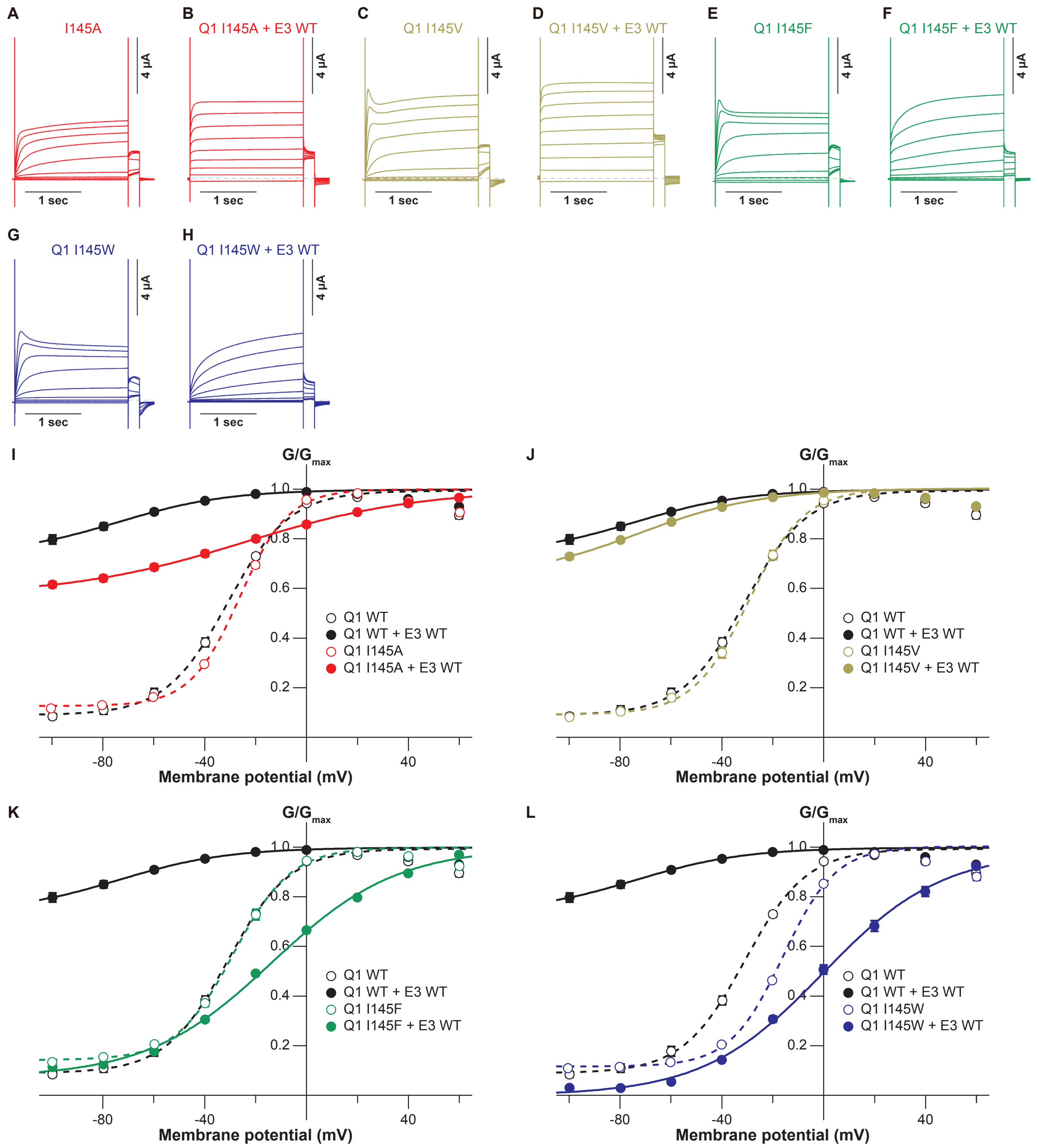

**Figure supplement 7, related to Figure 2. Current traces and G-V relationships of KCNQ1 I145 mutants.**

**(A-L)** Representative current traces (A-H) and G-V relationships (I-L) of KCNQ1 I145 mutants with or without KCNE3 WT. Error bars indicate ± s.e.m. for $n$ = 10 in (I-L).

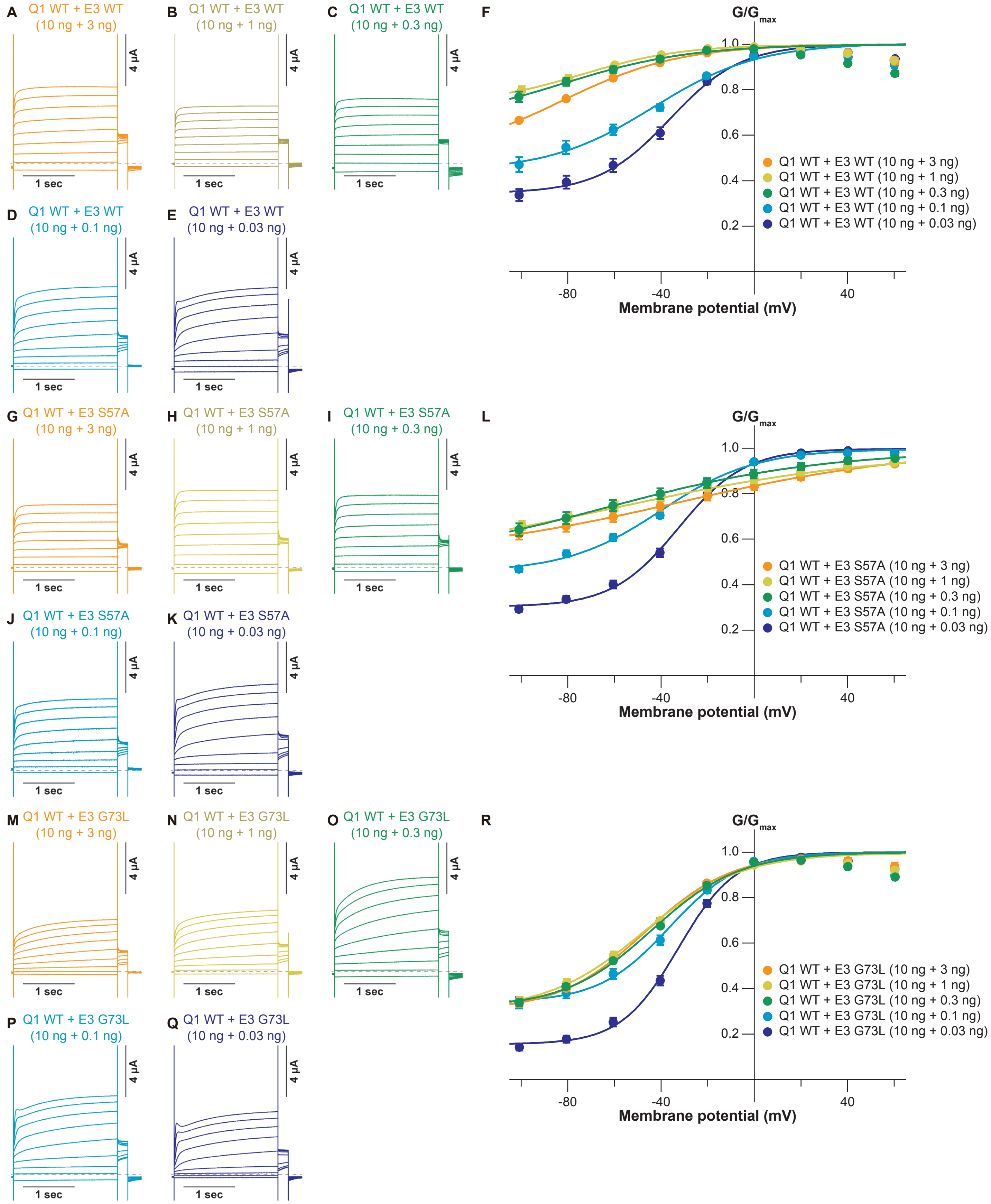

**Figure supplement 8, related to Figure 3. Dose-dependent modulation of KCNQ1 by decreasing amounts of KCNE3 WT and mutants.** (**A-F**) Representative current traces (A-E) and G-V relationships (B) of KCNQ1 WT co-expressed with each amount of KCNE3 WT. (**G-L**) Representative current traces (G-K) and G-V relationships (L) of KCNQ1 WT co-expressed with each amount of the KCNE3 S57A mutant. (**M-R**) Representative current traces (M-Q) and G-V relationships (F) of KCNQ1 WT co-expressed with each amount of the KCNE3 G73L mutant. Error bars indicate ± s.e.m. for $n$ = 10 in 10 ng KCNQ1 with 1 ng KCNE3 conditions and for $n$ = 5 in the other conditions in (F,L,R). For clear comparisons, the current traces and G-V relationships of 10 ng KCNQ1 WT with 1 ng KCNE3 WT and mutants shown in Figures 2B, 3B and 3J are redisplayed here.

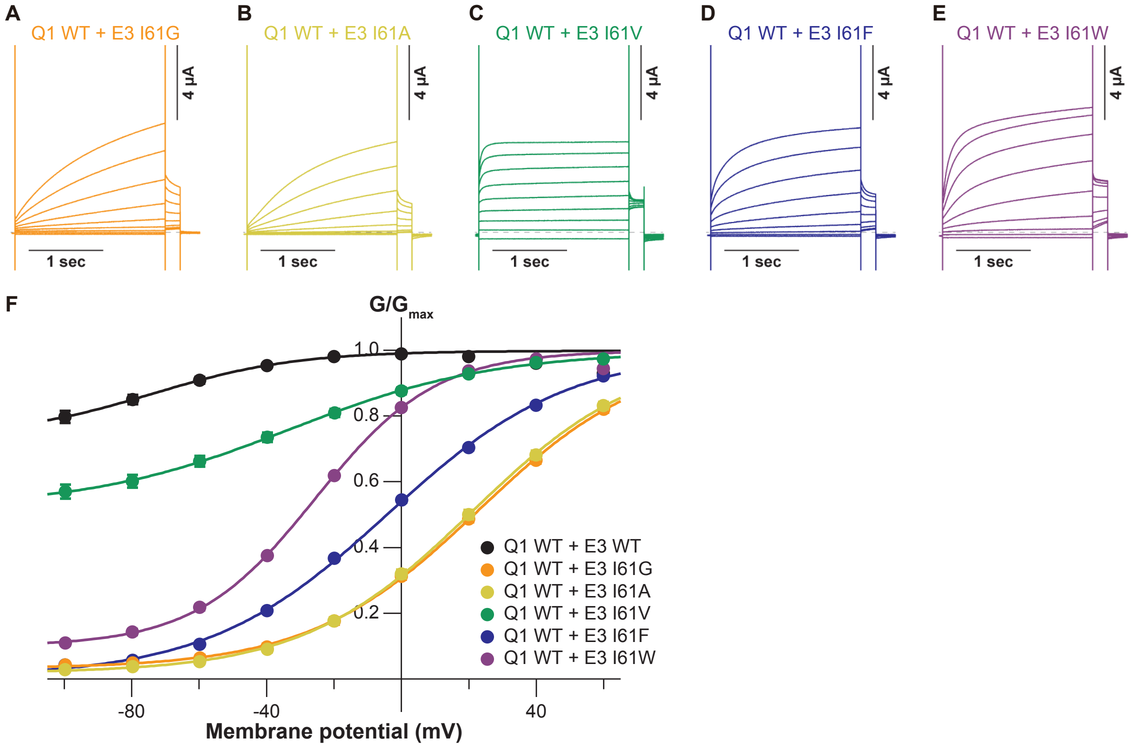

**Figure supplement 9, related to Figure 3. Current traces and G-V relationships of KCNQ1 WT with the KCNE3 I61 mutants.** (**A-F**) Representative current traces (A-E) and G-V relationships (F) of KCNQ1 WT with the KCNE3 I61 mutants. Error bars indicate ± s.e.m. for $n$ = 10 in (F).

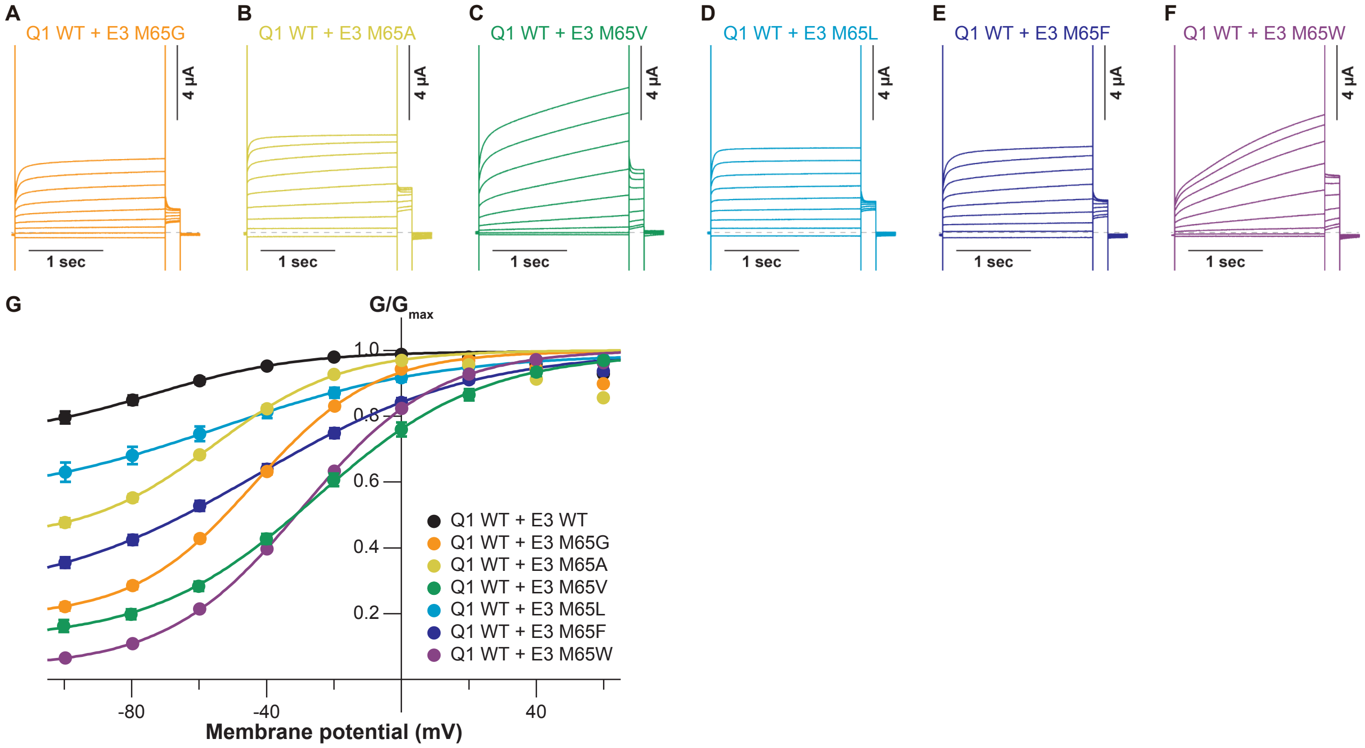

**Figure supplement 10, related to Figure 3. Current traces and G-V relationships of KCNQ1 WT with the KCNE3 M65 mutants.** (**A-F**) Representative current traces (A-E) and G-V relationships (F) of KCNQ1 WT with the KCNE3 M65 mutants. Error bars indicate ± s.e.m. for *n* = 10 in (F).

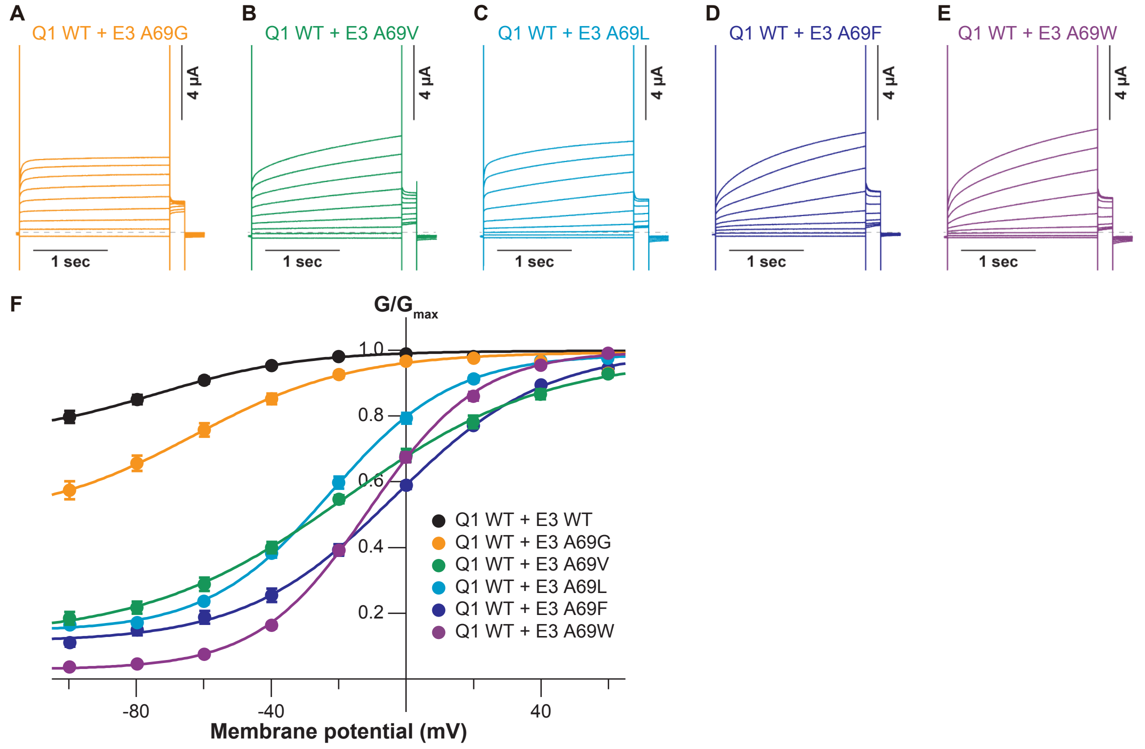

**Figure supplement 11, related to Figure 3. Current traces and G-V relationships of KCNQ1 WT with the KCNE3 A69 mutants.**

(**A-F**) Representative current traces (A-E) and G-V relationships (F) of KCNQ1 WT with the KCNE3 A69 mutants. Error bars indicate ± s.e.m. for $n$ = 10 in (F).

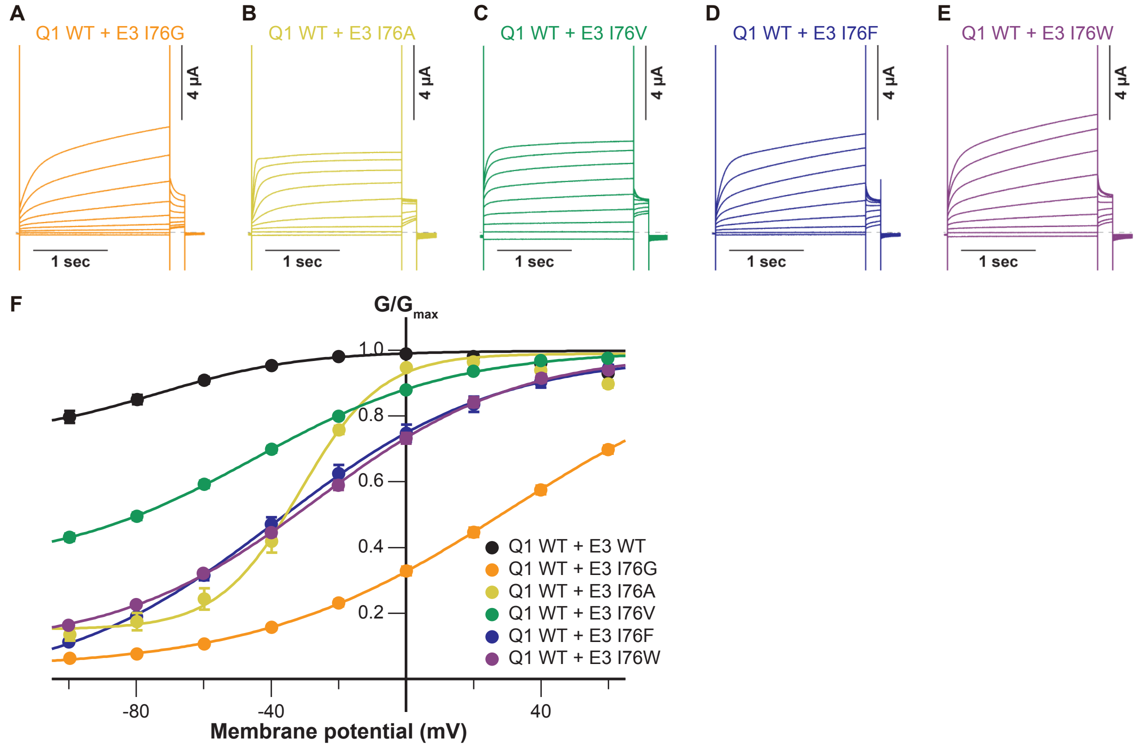

**Figure supplement 12, related to Figure 3. Current traces and G-V relationships of KCNQ1 WT with the KCNE3 I76 mutants.**
(**A-F**) Representative current traces (A-E) and G-V relationships (F) of KCNQ1 WT with the KCNE3 I76 mutants. Error bars indicate ± s.e.m. for *n* = 10 in (F).

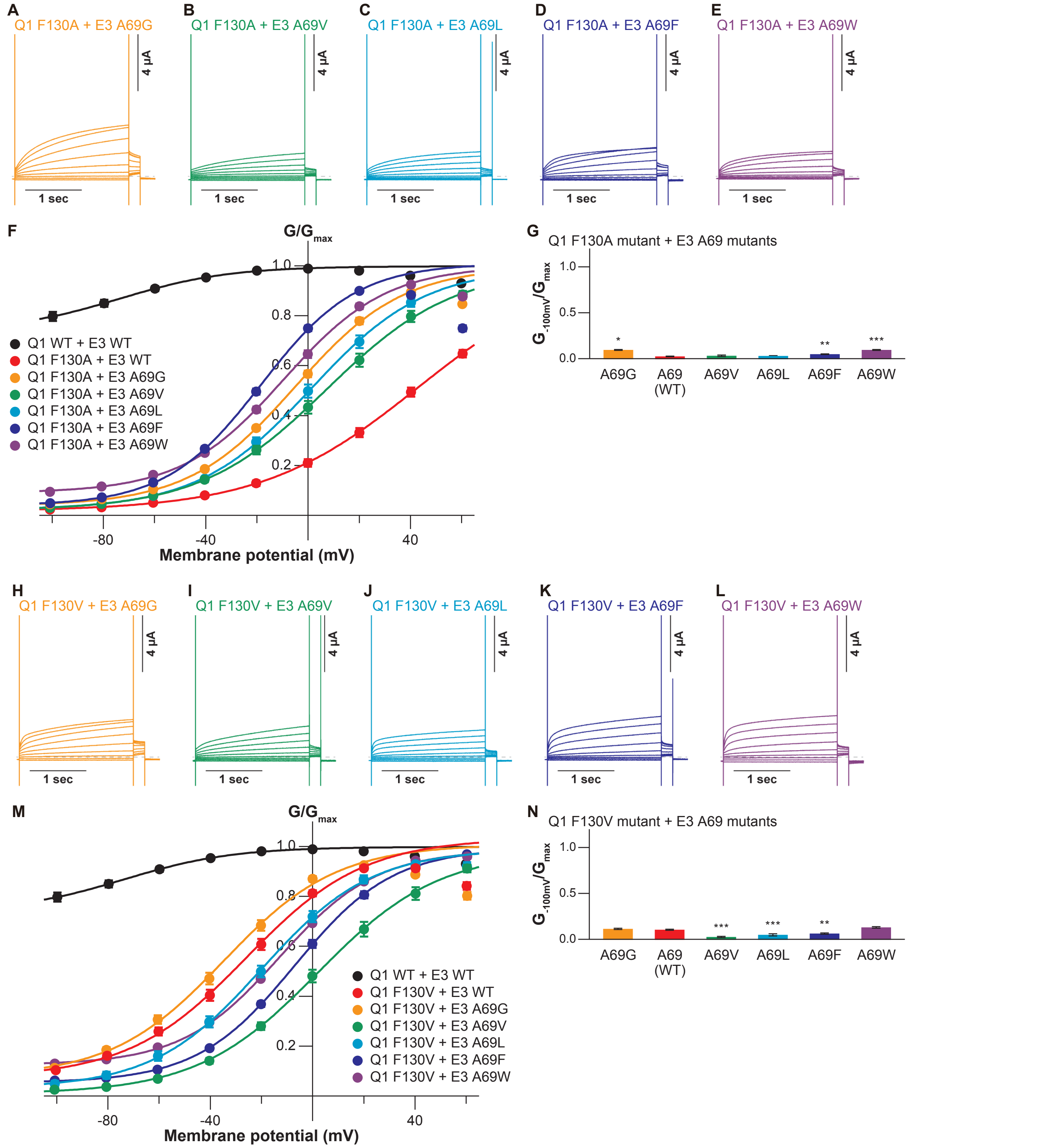

**Figure supplement 13, related to Figure 4. Current traces, G-V relationships, and ratios of conductance of the KCNQ1 F130 mutants with the KCNE3 A69 mutants.**

(**A-G**) Representative current traces (A-E), G-V relationships (F), and ratios of conductance at -100 mV ($G_{-100mV}$) and maximum conductance ($G_{max}$) (G) of the KCNQ1 F130A mutant with the KCNE3 A69 mutants. (**H-N**) Representative current traces (H-L), G-V relationships (M), and ratios of conductance at -100 mV ($G_{-100mV}$) and maximum conductance ($G_{max}$) (N) of the KCNQ1 F130V mutant with the KCNE3 A69 mutants. Error bars indicate ± s.e.m. for *n* = 10 in (F,G,M,N).

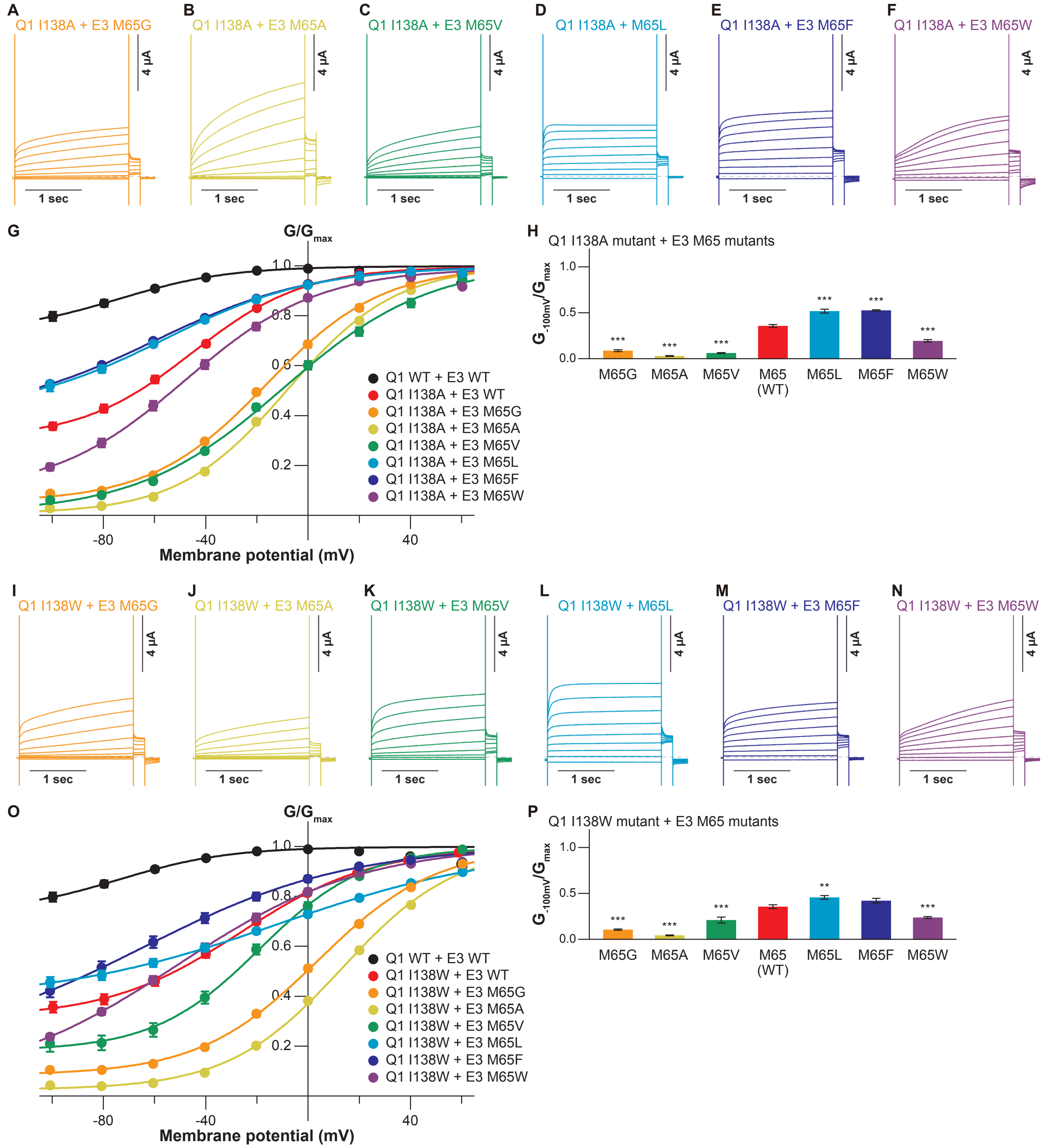

**Figure supplement 14, related to Figure 4. Current traces, G-V relationships, and ratios of conductance of the KCNQ1 I138 mutants with the KCNE3 M65 mutants.**

(**A-H**) Representative current traces (A-F), G-V relationships (G), and ratios of conductance at -100 mV ($G_{-100mV}$) and maximum conductance ($G_{max}$) (H) of the KCNQ1 I138A mutant with the KCNE3 M65 mutants. (**I-P**) Representative current traces (I-N), G-V relationships (O), and ratios of conductance at -100 mV ($G_{-100mV}$) and maximum conductance ($G_{max}$) (P) of the KCNQ1 I138W mutant with the KCNE3 M65 mutants. Error bars indicate ± s.e.m. for *n* = 10 in (G,H,O,P).

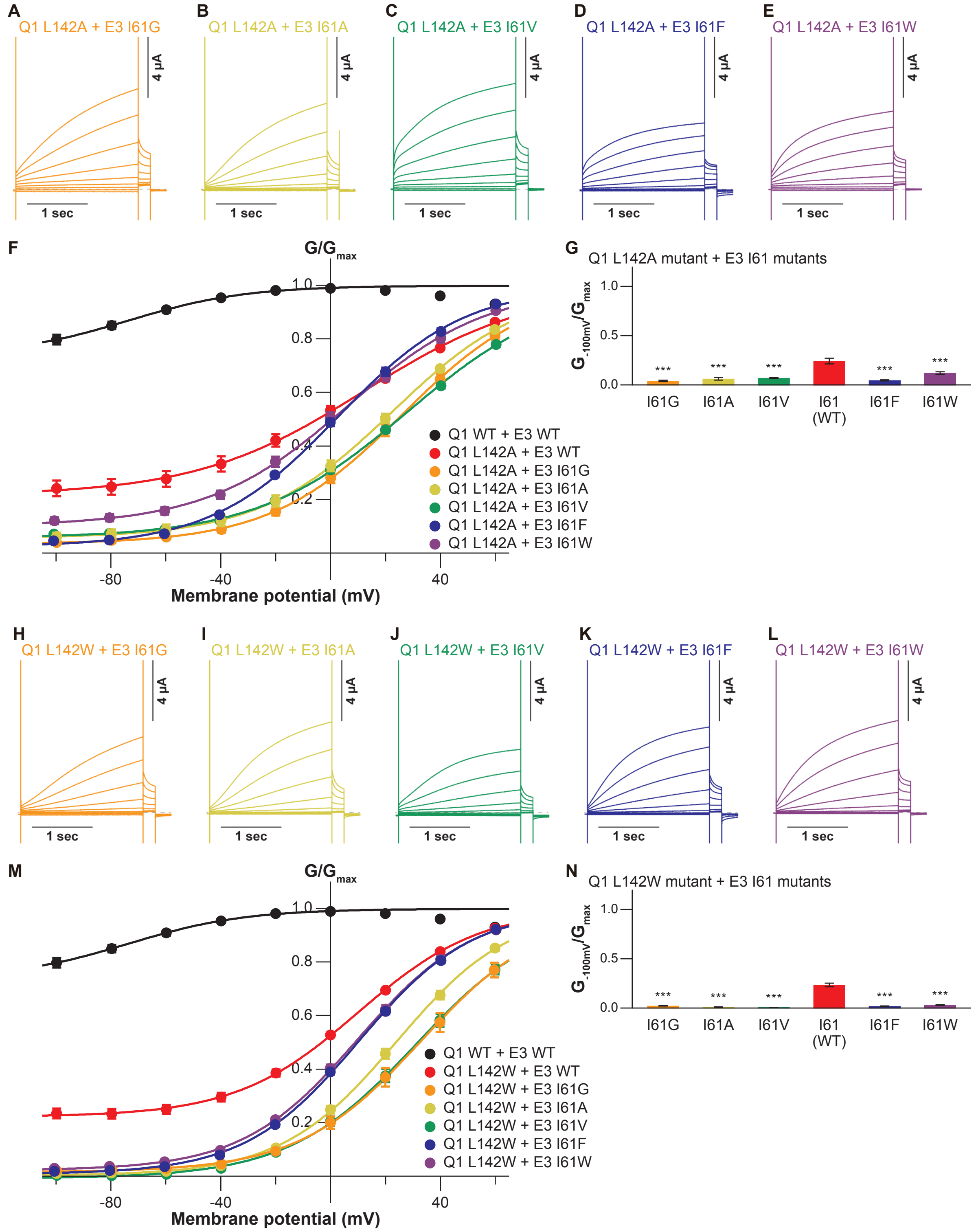

**Figure supplement 15, related to Figure 4. Current traces, G-V relationships, and ratios of conductance of the KCNQ1 L142 mutants with the KCNE3 I61 mutants.**

(**A-G**) Representative current traces (A-E), G-V relationships (F), and ratios of conductance at -100 mV ($G_{-100mV}$) and maximum conductance ($G_{max}$) (G) of the KCNQ1 L142A mutant with the KCNE3 I61 mutants. (**H-N**) Representative current traces (H-L), G-V relationships (M), and ratios of conductance at -100 mV ($G_{-100mV}$) and maximum conductance ($G_{max}$) (N) of the KCNQ1 L142W mutant with the KCNE3 I61 mutants. Error bars indicate ± s.e.m. for $n$ = 10 in (F,G,M,M).

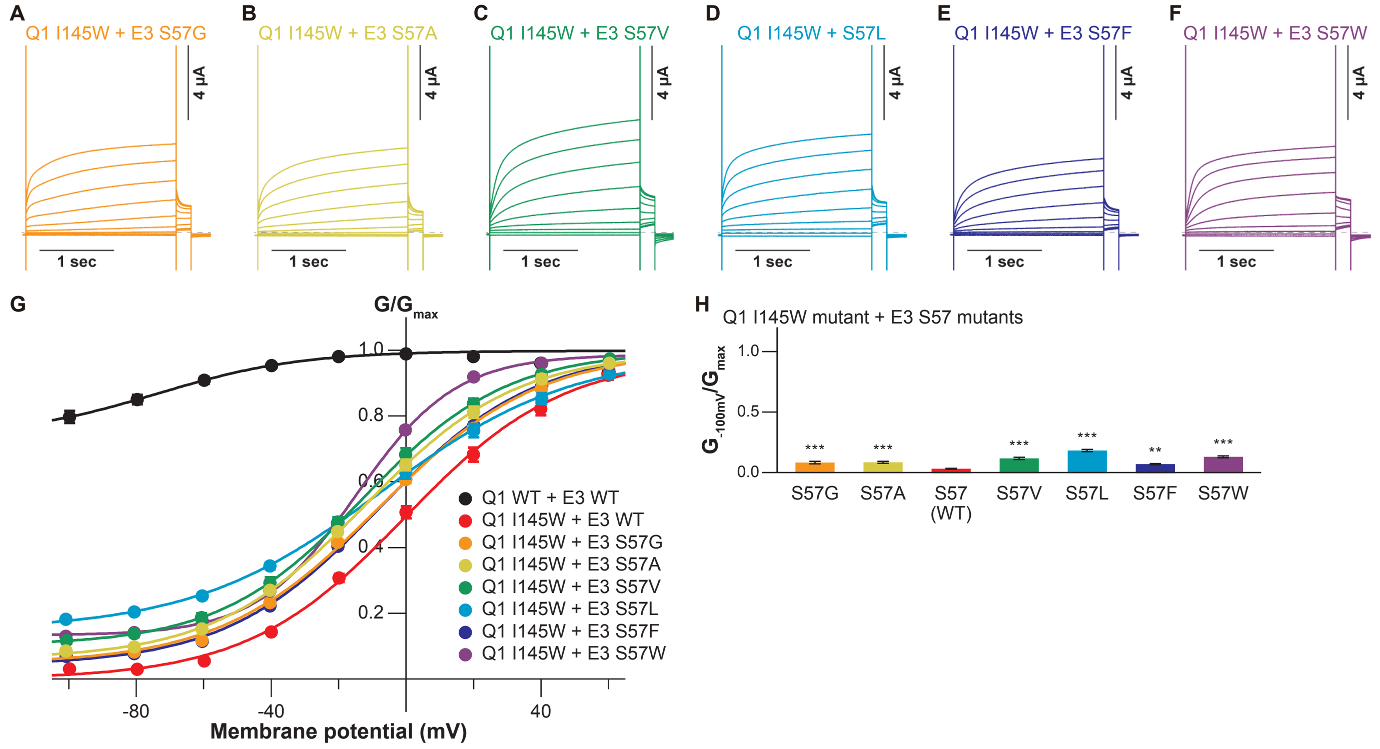

**Figure supplement 16, related to Figure 4. Current traces, G-V relationships, and ratios of conductance of the KCNQ1 I145W mutant with the KCNE3 S57 mutants.**

(**A-H**) Representative current traces (A-F), G-V relationships (G), and ratios of conductance at -100 mV ($G_{-100mV}$) and maximum conductance ($G_{max}$) (H) of the KCNQ1 I145W mutant with the KCNE3 S57 mutants. Error bars indicate ± s.e.m. for n = 10 in (G,H).

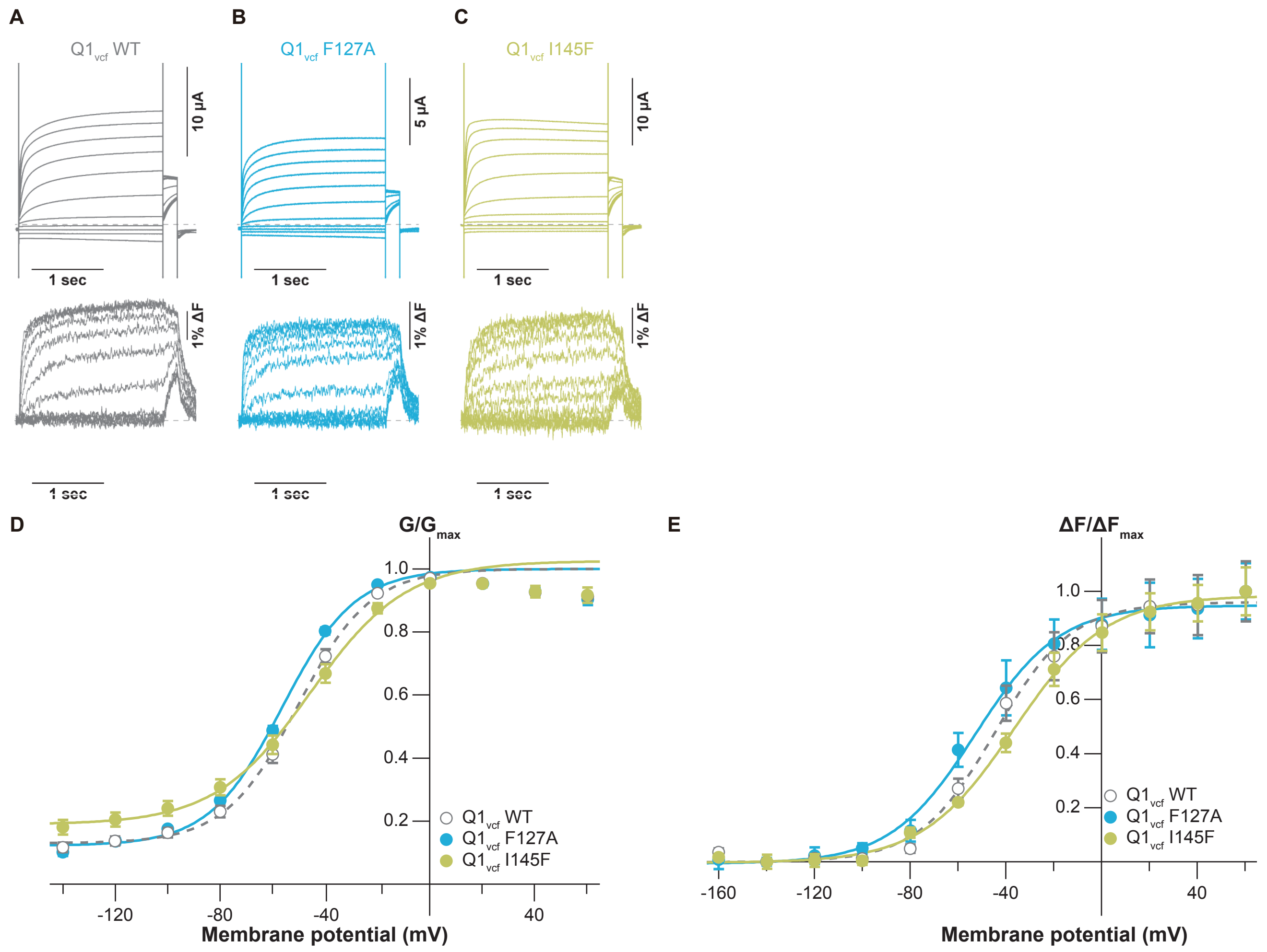

**Figure supplement 17, related to Figure 5. Conductance-voltage and fluorescence–voltage relationships for KCNQ1 mutants.** (**A-C**) Ionic currents (upper row) and fluorescence traces (lower row) of $KCNQ1_{vcf}$ WT (A), $KCNQ1_{vcf}$ F127A (B), and $KCNQ1_{vcf}$ I145F (C). (**D,E**) G-V (D) and F-V (E) relationships of $KCNQ1_{vcf}$ WT, $KCNQ1_{vcf}$ F127A, and $KCNQ1_{vcf}$ I145F. Error bars indicate ± s.e.m. for $n = 5$ in (D,E).

| Conditions | $I_{max}$ (µA) | $V_{1/2}$ (mV) | $z$ | $G_{-100mV}/G_{max}$ | $n$ |
|---|---|---|---|---|---|
| KCNQ1 WT | 1.7± 0.2 | -31.0 ± 0.6 | 2.0 ± 0.0 | 0.08 ± 0.01 | 10 |
| KCNQ1 WT + KCNE3 WT (10 ng + 3 ng) | 2.3 ± 0.3 | n.d. | n.d. | 0.67 ± 0.01 | 5 |
| KCNQ1 WT + KCNE3 WT (10 ng + 1 ng) | 2.2 ± 0.4 | n.d. | n.d. | 0.80 ± 0.02 | 10 |
| KCNQ1 WT + KCNE3 WT (10 ng + 0.3 ng) | 2.7 ± 0.4 | n.d. | n.d. | 0.77 ± 0.02 | 5 |
| KCNQ1 WT + KCNE3 WT (10 ng + 0.1 ng) | 2.8 ± 0.4 | -43.4 ± 1.2 | 1.2 ± 0.0 | 0.47 ± 0.03 | 5 |
| KCNQ1 WT + KCNE3 WT (10 ng + 0.03 ng) | 2.5 ± 0.2 | -36.3 ± 1.2 | 1.6 ± 0.1 | 0.34 ± 0.03 | 5 |
| KCNQ1 F123A | 1.5± 0.1 | -31.9 ± 0.6 | 2.2 ± 0.0 | 0.12 ± 0.01 | 10 |
| KCNQ1 F123A + KCNE3 WT | 1.3± 0.1 | n.d. | n.d. | 0.61 ± 0.01 | 10 |
| KCNQ1 F123W | 1.4± 0.3 | -37.8 ± 0.3 | 2.3 ± 0.0 | 0.07 ± 0.00 | 10 |
| KCNQ1 F123W + KCNE3 WT | 2.3± 0.2 | n.d. | n.d. | 0.66 ± 0.01 | 10 |
| KCNQ1 F127A | 1.4 ± 0.2 | -34.4 ± 0.6 | 2.2 ± 0.1 | 0.11 ± 0.01 | 10 |
| KCNQ1 F127A + KCNE3 WT | 1.6 ± 0.2 | -29.0 ± 0.5 | 1.3 ± 0.0 | 0.17 ± 0.02 | 10 |
| KCNQ1 F127V | 1.1 ± 0.1 | -27.0 ± 0.5 | 2.5 ± 0.0 | 0.09 ± 0.01 | 10 |
| KCNQ1 F127V + KCNE3 WT | 1.7 ± 0.2 | -41.3 ± 0.4 | 1.4 ± 0.0 | 0.25 ± 0.02 | 10 |
| KCNQ1 F127L | 1.4 ± 0.1 | -30.1 ± 0.2 | 2.1 ± 0.0 | 0.11 ± 0.01 | 10 |
| KCNQ1 F127L + KCNE3 WT | 1.7 ± 0.2 | -29.8 ± 1.0 | 1.4 ± 0.0 | 0.38 ± 0.02 | 10 |
| KCNQ1 F127W | 2.3 ± 0.2 | -26.7 ± 0.4 | 2.6 ± 0.1 | 0.04 ± 0.00 | 10 |
| KCNQ1 F127W + KCNE3 WT | 1.2 ± 0.1 | n.d. | n.d. | 0.61 ± 0.01 | 10 |
| KCNQ1 F130A | 1.1 ± 0.1 | -23.9 ± 0.7 | 2.5 ± 0.0 | 0.06 ± 0.00 | 10 |
| KCNQ1 F130A + KCNE3 WT | 1.1 ± 0.1 | 42.3 ± 2.1 | 0.8 ± 0.0 | 0.02 ± 0.00 | 10 |
| KCNQ1 F130V | 1.3 ± 0.0 | -16.6 ± 1.1 | 2.3 ± 0.1 | 0.05 ± 0.01 | 10 |
| KCNQ1 F130V + KCNE3 WT | 0.9 ± 0.1 | -27.8 ± 2.1 | 1.2 ± 0.0 | 0.10 ± 0.01 | 10 |
| KCNQ1 F130L | 1.7 ± 0.2 | -20.7 ± 0.4 | 2.1 ± 0.0 | 0.06 ± 0.01 | 10 |
| KCNQ1 F130L + KCNE3 WT | 1.5 ± 0.2 | 10.3 ± 2.2 | 1.1 ± 0.0 | 0.01 ± 0.00 | 10 |
| KCNQ1 F130W | 1.2 ± 0.1 | -20.3 ± 0.4 | 2.1 ± 0.0 | 0.25 ± 0.02 | 10 |
| KCNQ1 F130W + KCNE3 WT | 2.1 ± 0.2 | -9.8 ± 0.8 | 0.8 ± 0.0 | 0.37 ± 0.02 | 10 |
| KCNQ1 L134A | 1.7 ± 0.3 | -26.9 ± 0.6 | 2.3 ± 0.1 | 0.07 ± 0.01 | 10 |
| KCNQ1 L134A + KCNE3 WT | 1.8 ± 0.2 | -42.2 ± 1.4 | 1.0 ± 0.0 | 0.25 ± 0.01 | 10 |
| KCNQ1 L134V | 1.1 ± 0.2 | -27.4 ± 0.2 | 2.2 ± 0.0 | 0.04 ± 0.00 | 10 |
| KCNQ1 L134V + KCNE3 WT | 1.8 ± 0.2 | -18.4 ± 0.8 | 1.8 ± 0.1 | 0.08 ± 0.01 | 10 |
| KCNQ1 L134F | 1.1 ± 0.1 | n.d. | n.d. | 0.31 ± 0.02 | 10 |
| KCNQ1 L134F + KCNE3 WT | 1.5 ± 0.1 | -25.4 ± 1.1 | 1.3 ± 0.0 | 0.35 ± 0.02 | 10 |
| KCNQ1 L134W | 0.4 ± 0.0 | -31.1 ± 1.6 | 1.4 ± 0.1 | 0.10 ± 0.02 | 10 |
| KCNQ1 L134W + KCNE3 WT | 0.6 ± 0.1 | n.d. | n.d. | 0.68 ± 0.01 | 10 |
| KCNQ1 I138A | 2.2 ± 0.4 | -29.0 ± 0.4 | 2.1 ± 0.0 | 0.05 ± 0.00 | 10 |
| KCNQ1 I138A + KCNE3 WT | 2.6 ± 0.2 | -45.5 ± 0.9 | 1.1 ± 0.0 | 0.36 ± 0.02 | 10 |
| KCNQ1 I138V | 1.2 ± 0.1 | -31.2 ± 0.5 | 2.1 ± 0.0 | 0.07 ± 0.01 | 10 |
| KCNQ1 I138V + KCNE3 WT | 1.6 ± 0.1 | n.d. | n.d. | 0.66 ± 0.01 | 10 |
| KCNQ1 I138F | 1.1 ± 0.2 | -29.7 ± 0.4 | 2.2 ± 0.0 | 0.08 ± 0.01 | 10 |
| KCNQ1 I138F + KCNE3 WT | 2.2 ± 0.1 | n.d. | n.d. | 0.64 ± 0.01 | 10 |
| KCNQ1 I138W | 1.8 ± 0.3 | -28.1 ± 0.6 | 2.3 ± 0.0 | 0.13 ± 0.01 | 10 |
| KCNQ1 I138W + KCNE3 WT | 2.6 ± 0.3 | -26.2 ± 0.8 | 1.0 ± 0.0 | 0.36 ± 0.02 | 10 |
| KCNQ1 L142A | 1.5 ± 0.1 | -37.7 ± 0.4 | 2.7 ± 0.1 | 0.16 ± 0.00 | 10 |
| KCNQ1 L142A + KCNE3 WT | 1.3 ± 0.1 | 12.5 ± 1.1 | 0.8 ± 0.0 | 0.24 ± 0.03 | 10 |
| KCNQ1 L142V | 2.1 ± 0.2 | -35.2 ± 0.5 | 2.3 ± 0.1 | 0.10 ± 0.01 | 10 |
| KCNQ1 L142V + KCNE3 WT | 2.5 ± 0.2 | -31.5 ± 0.8 | 0.9 ± 0.0 | 0.42 ± 0.02 | 10 |
| KCNQ1 L142F | 1.1 ± 0.1 | -30.3 ± 0.5 | 2.4 ± 0.0 | 0.09 ± 0.00 | 10 |
| KCNQ1 L142F + KCNE3 WT | 2.4 ± 0.1 | n.d. | n.d. | 0.60 ± 0.01 | 10 |
| KCNQ1 L142W | 1.0 ± 0.1 | -32.0 ± 0.5 | 2.5 ± 0.0 | 0.10 ± 0.01 | 10 |
| KCNQ1 L142W + KCNE3 WT | 1.7 ± 0.1 | 10.1 ± 0.7 | 1.1 ± 0.0 | 0.23 ± 0.02 | 10 |
| KCNQ1 I145A | 1.7 ± 0.2 | -24.4 ± 0.3 | 2.6 ± 0.0 | 0.11 ± 0.01 | 10 |
| KCNQ1 I145A + KCNE3 WT | 1.7 ± 0.2 | n.d. | n.d. | 0.62 ± 0.01 | 10 |
| KCNQ1 I145V | 2.1 ± 0.2 | -29.6 ± 0.9 | 2.4 ± 0.1 | 0.08 ± 0.01 | 10 |
| KCNQ1 I145V + KCNE3 WT | 2.5 ± 0.1 | n.d. | n.d. | 0.73 ± 0.01 | 10 |
| KCNQ1 I145F | 1.3 ± 0.2 | -28.5 ± 1.0 | 2.3 ± 0.1 | 0.13 ± 0.01 | 10 |
| KCNQ1 I145F + KCNE3 WT | 1.7 ± 0.1 | -13.4 ± 0.7 | 1.0 ± 0.0 | 0.11 ± 0.01 | 10 |
| KCNQ1 I145W | 1.8 ± 0.1 | -16.1 ± 0.3 | 2.5 ± 0.1 | 0.11 ± 0.01 | 10 |
| KCNQ1 I145W + KCNE3 WT | 1.5 ± 0.2 | -0.1 ± 1.7 | 1.0 ± 0.1 | 0.03 ± 0.00 | 10 |

**Table supplement 1, related to Figure 2. Summary of the electrophysiological properties of KCNQ1 WT and mutants with or without KCNE3 WT.**

Maximum tail current amplitudes ($I_{max}$), parameters deduced from the Boltzmann fitting ($V_{1/2}$ and $z$), and ratios of conductance at -100 mV ($G_{-100mV}$) and maximum conductance ($G_{max}$) of KCNQ1 WT and mutants with or without KCNE3 WT. $n$ is the number of experiments.

| Conditions | $I_{max}$ (µA) | $V_{1/2}$ (mV) | $z$ | $G_{-100mV}/G_{max}$ | $n$ |
|---|---|---|---|---|---|
| KCNQ1 WT + KCNE3 S57G | 1.7 ± 0.3 | -26.5 ± 2.4 | 0.8 ± 0.1 | 0.32 ± 0.02 | 10 |
| KCNQ1 WT + KCNE3 S57A (10 ng + 3 ng) | 3.0 ± 0.5 | n.d. | n.d. | 0.62 ± 0.02 | 5 |
| KCNQ1 WT + KCNE3 S57A (10 ng + 1 ng) | 2.5 ± 0.2 | n.d. | n.d. | 0.66 ± 0.02 | 10 |
| KCNQ1 WT + KCNE3 S57A (10 ng + 0.3 ng) | 3.0 ± 0.3 | n.d. | n.d. | 0.64 ± 0.03 | 5 |
| KCNQ1 WT + KCNE3 S57A (10 ng + 0.1 ng) | 2.6 ± 0.3 | -40.3 ± 1.2 | 1.2 ± 0.1 | 0.47 ± 0.01 | 5 |
| KCNQ1 WT + KCNE3 S57A (10 ng + 0.03 ng) | 2.6 ± 0.3 | -32.4 ± 0.9 | 1.8 ± 0.1 | 0.29 ± 0.01 | 5 |
| KCNQ1 WT + KCNE3 S57V | 2.5 ± 0.1 | 3.4 ± 1.5 | 0.6 ± 0.0 | 0.47 ± 0.01 | 10 |
| KCNQ1 WT + KCNE3 S57L | 2.0 ± 0.1 | -9.1 ± 1.5 | 0.8 ± 0.0 | 0.20 ± 0.01 | 10 |
| KCNQ1 WT + KCNE3 S57F | 1.9 ± 0.2 | -21.4 ± 1.8 | 1.2 ± 0.0 | 0.10 ± 0.01 | 10 |
| KCNQ1 WT + KCNE3 S57W | 2.3 ± 0.1 | -12.9 ± 1.0 | 1.0 ± 0.0 | 0.10 ± 0.01 | 10 |
| KCNQ1 WT + KCNE3 I61G | 3.1 ± 0.3 | 23.2 ± 1.1 | 1.0 ± 0.0 | 0.04 ± 0.01 | 10 |
| KCNQ1 WT + KCNE3 I61A | 3.4 ± 0.4 | 21.1 ± 1.5 | 1.0 ± 0.0 | 0.03 ± 0.01 | 10 |
| KCNQ1 WT + KCNE3 I61V | 2.3 ± 0.3 | n.d. | n.d. | 0.57 ± 0.02 | 10 |
| KCNQ1 WT + KCNE3 I61F | 3.4 ± 0.2 | -3.9 ± 0.9 | 0.9 ± 0.0 | 0.04 ± 0.00 | 10 |
| KCNQ1 WT + KCNE3 I61W | 2.2 ± 0.2 | -25.4 ± 0.6 | 1.4 ± 0.0 | 0.11 ± 0.01 | 10 |
| KCNQ1 WT + KCNE3 M65G | 2.0 ± 0.2 | -43.8 ± 0.6 | 1.4 ± 0.0 | 0.22 ± 0.01 | 10 |
| KCNQ1 WT + KCNE3 M65A | 2.3 ± 0.2 | -55.6 ± 0.8 | 1.3 ± 0.0 | 0.48 ± 0.01 | 10 |
| KCNQ1 WT + KCNE3 M65V | 1.5 ± 0.1 | -24.6 ± 1.5 | 1.0 ± 0.0 | 0.16 ± 0.02 | 10 |
| KCNQ1 WT + KCNE3 M65L | 2.0 ± 0.2 | n.d. | n.d. | 0.63 ± 0.03 | 10 |
| KCNQ1 WT + KCNE3 M65F | 2.4 ± 0.3 | -42.2 ± 1.5 | 0.8 ± 0.0 | 0.36 ± 0.02 | 10 |
| KCNQ1 WT + KCNE3 M65W | 1.7 ± 0.2 | -29.7 ± 0.4 | 1.3 ± 0.0 | 0.07 ± 0.01 | 10 |
| KCNQ1 WT + KCNE3 A69G | 1.9 ± 0.2 | n.d. | n.d. | 0.57 ± 0.03 | 10 |
| KCNQ1 WT + KCNE3 A69V | 2.0 ± 0.2 | -16.5 ± 1.6 | 0.8 ± 0.1 | 0.19 ± 0.02 | 10 |
| KCNQ1 WT + KCNE3 A69L | 1.6 ± 0.2 | -21.7 ± 1.3 | 1.4 ± 0.0 | 0.17 ± 0.01 | 10 |
| KCNQ1 WT + KCNE3 A69F | 2.3 ± 0.2 | -3.6 ± 0.8 | 1.1 ± 0.0 | 0.11 ± 0.01 | 10 |
| KCNQ1 WT + KCNE3 A69W | 2.1 ± 0.2 | -10.9 ± 1.0 | 1.5 ± 0.1 | 0.04 ± 0.01 | 10 |
| KCNQ1 WT + KCNE3 G73A | 2.4 ± 0.2 | n.d. | n.d. | 0.77 ± 0.01 | 10 |
| KCNQ1 WT + KCNE3 G73V | 1.6 ± 0.1 | -42.7 ± 1.1 | 1.4 ± 0.0 | 0.22 ± 0.02 | 10 |
| KCNQ1 WT + KCNE3 G73L (10 ng + 3 ng) | 1.8 ± 0.2 | -46.8 ± 1.0 | 1.3 ± 0.1 | 0.34 ± 0.01 | 5 |
| KCNQ1 WT + KCNE3 G73L (10 ng + 1 ng) | 2.0 ± 0.2 | -47.6 ± 1.1 | 1.2 ± 0.0 | 0.35 ± 0.02 | 10 |
| KCNQ1 WT + KCNE3 G73L (10 ng + 0.3 ng) | 2.2 ± 0.6 | -44.1 ± 0.9 | 1.4 ± 0.1 | 0.34 ± 0.02 | 5 |
| KCNQ1 WT + KCNE3 G73L (10 ng + 0.1 ng) | 2.1 ± 0.2 | -36.5 ± 0.9 | 1.7 ± 0.1 | 0.34 ± 0.03 | 5 |
| KCNQ1 WT + KCNE3 G73L (10 ng + 0.03 ng) | 1.9 ± 0.1 | -32.7 ± 1.1 | 2.1 ± 0.1 | 0.14 ± 0.01 | 5 |
| KCNQ1 WT + KCNE3 G73F | 2.1 ± 0.2 | -43.0 ± 1.1 | 1.2 ± 0.0 | 0.43 ± 0.02 | 10 |
| KCNQ1 WT + KCNE3 G73W | 2.3 ± 0.2 | -42.4 ± 1.7 | 1.2 ± 0.0 | 0.24 ± 0.01 | 10 |
| KCNQ1 WT + KCNE3 I76G | 4.0 ± 0.3 | 30.6 ± 2.1 | 0.7 ± 0.0 | 0.06 ± 0.01 | 10 |
| KCNQ1 WT + KCNE3 I76A | 2.3 ± 0.3 | -31.9 ± 1.2 | 2.1 ± 0.1 | 0.14 ± 0.02 | 10 |
| KCNQ1 WT + KCNE3 I76V | 1.7 ± 0.3 | -45.0 ± 1.3 | 0.9 ± 0.0 | 0.43 ± 0.01 | 10 |
| KCNQ1 WT + KCNE3 I76F | 1.7 ± 0.2 | -36.3 ± 2.2 | 0.8 ± 0.0 | 0.11 ± 0.01 | 10 |
| KCNQ1 WT + KCNE3 I76W | 1.8 ± 0.2 | -26.0 ± 1.5 | 0.8 ± 0.0 | 0.16 ± 0.01 | 10 |

**Table supplement 2, related to Figure 3. Summary of the electrophysiological properties of KCNQ1 WT with KCNE3 mutants.** Maximum tail current amplitudes ($I_{max}$), parameters deduced from the Boltzmann fitting ($V_{1/2}$ and $z$), and ratios of conductance at -100 mV ($G_{-100mV}$) and maximum conductance ($G_{max}$) of KCNQ1 WT with KCNE3 mutants. $n$ is the number of experiments.

| Conditions | $I_{max}$ (µA) | $V_{1/2}$ (mV) | *z* | $G_{-100mV}/G_{max}$ | *n* |
|---|---|---|---|---|---|
| KCNQ1 F127A + KCNE3 G73A | 1.5 ± 0.1 | -31.5 ± 0.6 | 1.7 ± 0.0 | 0.10 ± 0.01 | 10 |
| KCNQ1 F127A + KCNE3 G73V | 2.1 ± 0.2 | -46.6 ± 0.8 | 1.4 ± 0.0 | 0.36 ± 0.02 | 10 |
| KCNQ1 F127A + KCNE3 G73L | 2.4 ± 0.3 | n.d. | n.d. | 0.67 ± 0.01 | 10 |
| KCNQ1 F127A + KCNE3 G73F | 2.1 ± 0.2 | -55.6 ± 1.2 | 1.3 ± 0.0 | 0.45 ± 0.02 | 10 |
| KCNQ1 F127A + KCNE3 G73W | 1.8 ± 0.2 | -46.2 ± 1.0 | 1.3 ± 0.0 | 0.38 ± 0.01 | 10 |
| KCNQ1 F130A + KCNE3 A69G | 0.8 ± 0.1 | -9.3 ± 1.2 | 1.3 ± 0.0 | 0.10 ± 0.01 | 10 |
| KCNQ1 F130A + KCNE3 A69V | 0.7 ± 0.1 | 8.3 ± 2.5 | 1.0 ± 0.0 | 0.03 ± 0.01 | 10 |
| KCNQ1 F130A + KCNE3 A69L | 0.7 ± 0.1 | 1.8 ± 2.2 | 1.2 ± 0.0 | 0.03 ± 0.00 | 10 |
| KCNQ1 F130A + KCNE3 A69F | 1.5 ± 0.1 | -18.8 ± 0.8 | 1.4 ± 0.0 | 0.05 ± 0.00 | 10 |
| KCNQ1 F130A + KCNE3 A69W | 0.8 ± 0.1 | -9.3 ± 1.2 | 1.3 ± 0.0 | 0.10 ± 0.01 | 10 |
| KCNQ1 F130V + KCNE3 A69G | 1.2 ± 0.2 | -35.2 ± 2.1 | 1.2 ± 0.0 | 0.11 ± 0.01 | 10 |
| KCNQ1 F130V + KCNE3 A69V | 0.7 ± 0.1 | 3.8 ± 2.2 | 1.1 ± 0.1 | 0.03 ± 0.01 | 10 |
| KCNQ1 F130V + KCNE3 A69L | 0.8 ± 0.1 | -19.3 ± 1.8 | 1.2 ± 0.1 | 0.05 ± 0.01 | 10 |
| KCNQ1 F130V + KCNE3 A69F | 1.1 ± 0.1 | -6.5 ± 1.3 | 1.3 ± 0.0 | 0.06 ± 0.01 | 10 |
| KCNQ1 F130V + KCNE3 A69W | 0.8 ± 0.1 | -12.3 ± 0.6 | 1.3 ± 0.0 | 0.13 ± 0.01 | 10 |
| KCNQ1 I138A + KCNE3 M65G | 1.6 ± 0.2 | -15.8 ± 1.2 | 1.1 ± 0.0 | 0.09 ± 0.01 | 10 |
| KCNQ1 I138A + KCNE3 M65A | 1.8 ± 0.2 | -8.0 ± 1.0 | 1.2 ± 0.0 | 0.03 ± 0.01 | 10 |
| KCNQ1 I138A + KCNE3 M65V | 1.7 ± 0.1 | -9.2 ± 1.5 | 0.9 ± 0.0 | 0.06 ± 0.01 | 10 |
| KCNQ1 I138A + KCNE3 M65L | 2.1 ± 0.3 | n.d. | n.d. | 0.52 ± 0.02 | 10 |
| KCNQ1 I138A + KCNE3 M65F | 1.6 ± 0.1 | n.d. | n.d. | 0.53 ± 0.01 | 10 |
| KCNQ1 I138A + KCNE3 M65W | 1.9 ± 0.2 | -47.9 ± 1.6 | 1.0 ± 0.0 | 0.19 ± 0.01 | 10 |
| KCNQ1 I138W + KCNE3 M65G | 1.5 ± 0.2 | 3.7 ± 1.1 | 1.1 ± 0.0 | 0.11 ± 0.01 | 10 |
| KCNQ1 I138W + KCNE3 M65A | 2.2 ± 0.3 | 13.5 ± 1.1 | 1.1 ± 0.0 | 0.04 ± 0.01 | 10 |
| KCNQ1 I138W + KCNE3 M65V | 2.6 ± 0.2 | -19.0 ± 0.9 | 1.2 ± 0.0 | 0.21 ± 0.03 | 10 |
| KCNQ1 I138W + KCNE3 M65L | 2.3 ± 0.3 | -10.9 ± 2.0 | 0.6 ± 0.1 | 0.46 ± 0.02 | 10 |
| KCNQ1 I138W + KCNE3 M65F | 2.2 ± 0.3 | -64.2 ± 2.7 | 0.7 ± 0.0 | 0.42 ± 0.03 | 10 |
| KCNQ1 I138W + KCNE3 M65W | 2.1 ± 0.3 | -50.3 ± 1.9 | 0.7 ± 0.0 | 0.24 ± 0.01 | 10 |
| KCNQ1 L142A + KCNE3 I61G | 3.8 ± 0.3 | 26.0 ± 2.0 | 1.1 ± 0.0 | 0.04 ± 0.01 | 10 |
| KCNQ1 L142A + KCNE3 I61A | 2.8 ± 0.2 | 22.3 ± 1.4 | 1.0 ± 0.0 | 0.06 ± 0.01 | 10 |
| KCNQ1 L142A + KCNE3 I61V | 3.8 ± 0.4 | 28.0 ± 1.1 | 1.0 ± 0.0 | 0.07 ± 0.01 | 10 |
| KCNQ1 L142A + KCNE3 I61F | 2.0 ± 0.1 | 3.2 ± 1.4 | 1.1 ± 0.0 | 0.05 ± 0.01 | 10 |
| KCNQ1 L142A + KCNE3 I61W | 2.3 ± 0.2 | 6.7 ± 2.2 | 1.0 ± 0.0 | 0.12 ± 0.01 | 10 |
| KCNQ1 L142W + KCNE3 I61G | 2.8 ± 0.2 | 33.5 ± 3.3 | 1.2 ± 0.0 | 0.02 ± 0.00 | 10 |
| KCNQ1 L142W + KCNE3 I61A | 3.1 ± 0.2 | 24.0 ± 1.5 | 1.2 ± 0.0 | 0.01 ± 0.00 | 10 |
| KCNQ1 L142W + KCNE3 I61V | 2.2 ± 0.2 | 31.5 ± 2.3 | 1.1 ± 0.0 | 0.00 ± 0.00 | 10 |
| KCNQ1 L142W + KCNE3 I61F | 2.8 ± 0.2 | 10.3 ± 0.9 | 1.2 ± 0.0 | 0.02 ± 0.00 | 10 |
| KCNQ1 L142W + KCNE3 I61W | 2.7 ± 0.2 | 10.0 ± 1.3 | 1.2 ± 0.0 | 0.03 ± 0.00 | 10 |
| KCNQ1 I145F + KCNE3 S57G | 2.0 ± 0.1 | 4.6 ± 1.3 | 0.9 ± 0.0 | 0.43 ± 0.01 | 10 |
| KCNQ1 I145F + KCNE3 S57A | 1.4 ± 0.2 | n.d. | n.d. | 0.61 ± 0.02 | 10 |
| KCNQ1 I145F + KCNE3 S57V | 2.0 ± 0.2 | 5.9 ± 0.7 | 0.9 ± 0.0 | 0.13 ± 0.01 | 10 |
| KCNQ1 I145F + KCNE3 S57L | 2.5 ± 0.2 | -7.8 ± 1.8 | 0.8 ± 0.0 | 0.16 ± 0.01 | 10 |
| KCNQ1 I145F + KCNE3 S57F | 1.9 ± 0.2 | -3.3 ± 1.3 | 0.9 ± 0.0 | 0.11 ± 0.01 | 10 |
| KCNQ1 I145F + KCNE3 S57W | 1.8 ± 0.2 | -14.2 ± 1.2 | 1.0 ± 0.0 | 0.09 ± 0.00 | 10 |
| KCNQ1 I145W + KCNE3 S57G | 2.4 ± 0.2 | -7.8 ± 1.2 | 1.1 ± 0.0 | 0.08 ± 0.01 | 10 |
| KCNQ1 I145W + KCNE3 S57A | 1.7 ± 0.2 | -11.8 ± 1.1 | 1.1 ± 0.1 | 0.09 ± 0.01 | 10 |
| KCNQ1 I145W + KCNE3 S57V | 1.7 ± 0.1 | -12.7 ± 1.8 | 1.2 ± 0.0 | 0.12 ± 0.01 | 10 |
| KCNQ1 I145W + KCNE3 S57L | 1.7 ± 0.1 | -6.3 ± 2.0 | 0.9 ± 0.0 | 0.18 ± 0.01 | 10 |
| KCNQ1 I145W + KCNE3 S57F | 2.0 ± 0.2 | -7.8 ± 1.0 | 1.1 ± 0.0 | 0.07 ± 0.01 | 10 |
| KCNQ1 I145W + KCNE3 S57W | 2.2 ± 0.3 | -14.2 ± 0.6 | 1.7 ± 0.0 | 0.13 ± 0.01 | 10 |

**Table supplement 3, related to Figure 4. Summary of the electrophysiological properties of KCNQ1 mutants with KCNE3 mutants.** Maximum tail current amplitudes ($I_{max}$), parameters deduced from the Boltzmann fitting ($V_{1/2}$ and $z$), and ratios of conductance at -100 mV ($G_{-100mV}$) and maximum conductance ($G_{max}$) of KCNQ1 mutants with KCNE3 mutants. $n$ is the number of experiments.

| Conditions | $I_{max}$ (µA) | $V_{1/2}$ (mV) | $z$ | $n$ |
|---|---|---|---|---|
| $KCNQ1_{vcf}$ WT | 4.8 ± 0.6 | -50.4 ± 1.4 | 1.8 ± 0.0 | 5 |
| $KCNQ1_{vcf}$ WT + KCNE3 WT | 16.9 ± 2.5 | n.d. | n.d. | 5 |
| $KCNQ1_{vcf}$ F127A | 2.3 ± 0.2 | -56.2 ± 1.0 | 1.8 ± 0.0 | 5 |
| $KCNQ1_{vcf}$ F127A + KCNE3 WT | 4.9 ± 0.6 | -96.8 ± 4.4 | 0.9 ± 0.0 | 5 |
| $KCNQ1_{vcf}$ F127A + KCNE3 G73L | 7.5 ± 1.6 | n.d. | n.d. | 5 |
| $KCNQ1_{vcf}$ I145F | 5.9 ± 0.8 | -46.8 ± 1.9 | 1.5 ± 0.0 | 5 |
| $KCNQ1_{vcf}$ I145F + KCNE3 WT | 4.8 ± 0.6 | -104.2 ± 8.5 | 0.5 ± 0.0 | 5 |
| $KCNQ1_{vcf}$ I145F + KCNE3 S57A | 9.0 ± 1.9 | n.d. | n.d. | 5 |

**Table supplement 4, related to Figure 5. Summary of the electrophysiological properties of $KCNQ1_{vcf}$ WT and mutants with or without KCNE3 mutants.**

Maximum tail current amplitudes ($I_{max}$) and parameters deduced from the Boltzmann fitting ($V_{1/2}$ and $z$) of $KCNQ1_{vcf}$ WT and mutants with or without KCNE3 mutants. $n$ is the number of experiments.

| **Conditions** | $V_{1/2(F1)}$ (mV) | $V_{1/2(F2)}$ (mV) | *n* |
|---|---|---|---|
| $KCNQ1_{vcf}$ WT | -44.3 ± 0.7 | n.d. | 5 |
| $KCNQ1_{vcf}$ WT + KCNE3 WT | -141.7 ± 10.8 | 96.2 ± 8.4 | 5 |
| $KCNQ1_{vcf}$ F127A | -52.3 ± 2.7 | n.d. | 5 |
| $KCNQ1_{vcf}$ F127A + KCNE3 WT | -77.1 ± 3.0 | 214.2 ± 20.2 | 5 |
| $KCNQ1_{vcf}$ F127A + KCNE3 G73L | -175.3 ± 27.3 | -41.8 ± 5.9 | 5 |
| $KCNQ1_{vcf}$ I145F | -36.6 ± 1.5 | n.d. | 5 |
| $KCNQ1_{vcf}$ I145F + KCNE3 WT | -121.2 ± 11.6 | 229.6 ± 25.3 | 5 |
| $KCNQ1_{vcf}$ I145F + KCNE3 S57A | -173.8 ± 11.2 | 238.1 ± 21.7 | 5 |

**Table supplement 5, related to Figure 5. Summary of the properties of $KCNQ1_{vcf}$ WT and mutants with or without KCNE3 mutants acquired from VCF recordings.**

Parameters deduced from the double Boltzmann fitting ($V_{1/2(F1)}$ and $V_{1/2(F2)}$) of $KCNQ1_{vcf}$ WT and mutants with or without KCNE3 mutants. *n* is the number of experiments.
